# Phenotypic Antibiotic Susceptibility Testing at the limit of one bacterial cell

**DOI:** 10.1101/2025.04.13.648565

**Authors:** Lisa Johansson, Irfan Ahmad, Jimmy Larsson, Spartak Zikrin, Petter Knagge, David Fange, Johan Elf

## Abstract

The blood of a patient with bacteremia contains only a few bacteria per milliliter. Recently, techniques have been developed for extracting these few bacteria from whole blood. Here, we evaluate if a phenotypic Antibiotic Susceptibility Test (AST) can be performed on a single bacterial cell by monitoring changes in its phenotypic distributions sampled over time rather than across populations, while also accounting for experimental noise and cell-to-cell variability. We use the method to estimate the time required to correctly classify single susceptible or resistant bacteria when treated with antibiotics that affect growth, cell division, or cause lysis. We find that due to heterogeneous single cell responses, it is necessary to consider all bacteria close to the susceptibility breakpoint as resistant. However, it is possible to follow up resistance calls by sequentially exposing the cell or its lineage to new antibiotics until a susceptible phenotype is detected.

## Introduction

Every year, 10 million people die from sepsis (Rudd et al. 2020). 1.5 million of these are associated with antibiotic resistance (Murray et al. 2022). The initial choice of antibiotic (AB) treatment is based on the experience and judgment of the treating physician (empirical treatment) because Antibiotic Susceptibility Tests (ASTs) are too slow. Current phenotypic ASTs (pASTs) require a positive blood culture (PBC) as the starting point. Here, a PBC means that patient blood has been mixed with broth and cultured until there are ≈10^8^ colony-forming units (CFU)/ml (Azrad et al. 2019). It typically takes 12-24h for the bacteria to reach this density since the bacterial viable count in the bloodstream is very low (<10 CFU/ml) (Wain et al. 1998; Kreger et al. 1980). The AST and species identification (ID) possibilities following PBCs have improved radically in the last five years. Phenotypic AST (pAST), i.e. AST methods based on how the bacteria respond to the drug, used to depend on plating assays, with a typical two-day turnaround time, including isolation streak from PBC. Now pASTs can be completed in 6-8 hours with the rapid EUCAST protocols (Jonasson et al. 2020). In addition, companies such as Q-linea, Gradientech and QuantaMatrix market assays that can perform pAST from a PBC in 4-6 hours (Reszetnik et al. 2024). The bottleneck for making a pAST that could guide antibiotic treatment in a few hours is the time to obtain a PBC. To remove this bottleneck, we need to determine antibiotic susceptibility directly on the handful of cells extracted from the patient’s blood instead of waiting for a PBC. The problem of extracting a single bacterium directly from a patient’s blood sample has been addressed in other papers, for example, (Kim et al. 2024; Marino Miguélez et al. 2025; Forsyth et al. 2021).

In (Baltekin et al. 2017), we showed that it’s possible to determine if a bacterial isolate is susceptible to an antibiotic in less than one generation by: (1) Monitoring the growth rate impact by the length extension of cells rather than measuring growth by an increase in the number of cells; (2) Averaging the growth rate over several (≈1000) bacteria to eliminate the measurement noise and biological cell-to-cell variation; (3) Normalizing the antibiotic impact to an untreated reference population of the same isolate to eliminate isolate-to-isolate variation. The question was whether we could perform pAST in less than 30 minutes, and the number of bacteria was not a critical concern. A product based on this assay has been shown to impact how antibiotics are prescribed for urinary tract infection (Martínez-Berganza Asensio et al. 2026).

Here, instead, the question is whether it is possible to perform pAST starting from a single bacterial cell, and if this is the case, how long it would take. The approach is to compare temporal changes in the distributions of the cell’s phenotypic properties before and after antibiotic treatment. This raises the question of whether the cell-to-cell variation in antibiotic response overshadows the difference between isolates with different susceptibility. The results from Brandis et al (Brandis et al. 2023) suggest that *E. coli* strains with an 8-fold difference in Minimum inhibitory concentrations (MICs) are well-separated on the single cell level for the cases of antibiotics that affect the growth rate, *i*.*e*. the rate of cell size expansion. However, for other species or antibiotics where the response is filamentation or lysis, the possibility of susceptibility calling on the single cell level is more uncertain. AST based on morphological analysis of bacteria in response to antibiotics was previously described by (Choi et al. 2014) where changes in average morphological features of bacteria were successfully used to determine antibiotic susceptibility. The susceptibility test was, however, carried out using information from tens of cells. (Zagajewski et al. 2023) showed that antibiotics that impact the chromosome organization in susceptible cells can be used to categorize individual fixed and stained *E. coli* as susceptible or resistant. This method is, however, not applicable in the single cell regime, since only a single antibiotic can be applied before irreversible fixation.

We will here investigate the feasibility of the suggested generic method for single-cell phenotypic AST based growth analysis of individual bacteria before and after treating the cell with an antibiotic. We propose the following consecutive steps. (i) monitor the cell before antibiotic treatment to establish a phenotypic baseline for the untreated cell, (ii) expose the cell to a first antibiotic and monitor the temporal phenotypic response in growth rate, or cell size and lysis, (iii) if necessary, repeated testing of other antibiotics, or concentrations, until an effective treatment is found. We will refer to the method as SPARTA-Single-cell Phenotypic Antibiotic Response Testing Assay.

The experiments are facilitated by growing and imaging bacteria in a microfluidic chip (Baltekin et al. 2017; Wang et al. 2010; Wallden et al. 2016), where the bacteria are imaged in time-lapse phase contrast microscopy (Figure 1a,d) and segmented using an AI model (methods). The segmented cells are tracked, and the time evolution of the area enclosed by the segment outline, referred to as the cell area (Figure 1b,e), is used to calculate growth rates for single cell lineages (Figure 1c,f).

**Figure 1.**
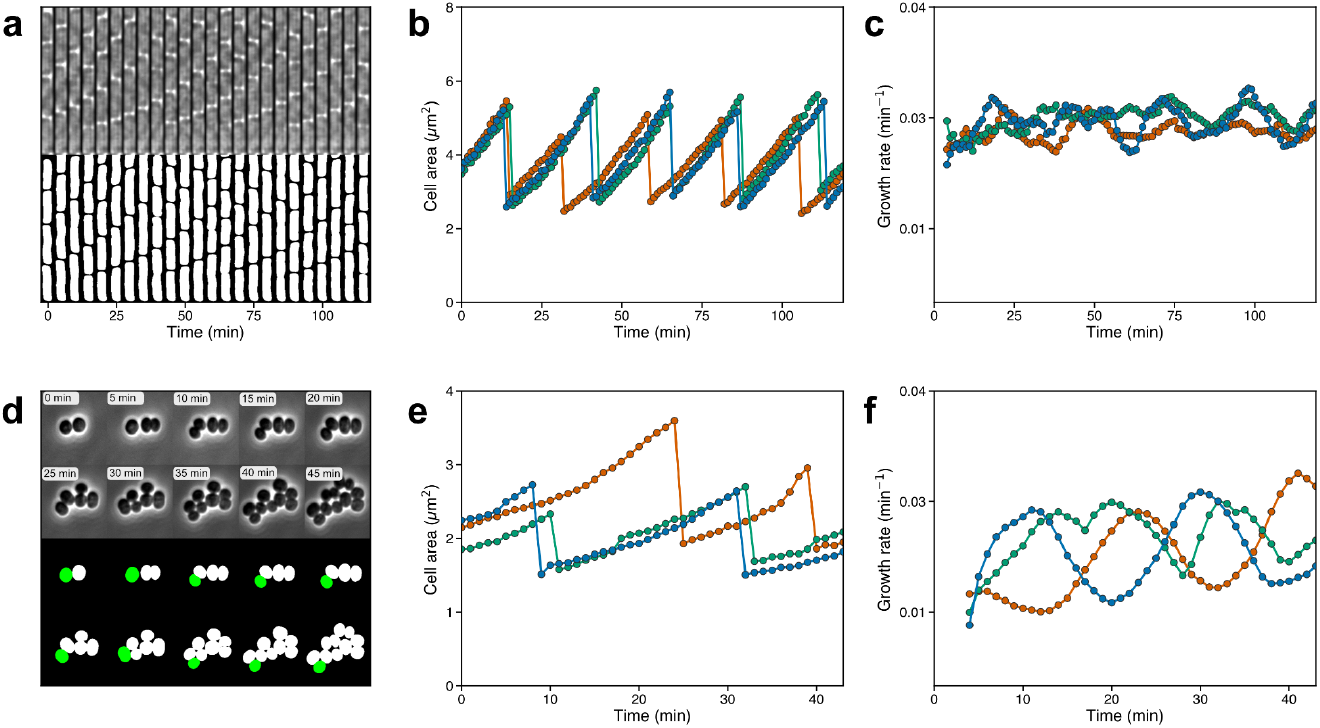
Illustration of single-cell time-lapse imaging and tracking. (**a, top**) Example of time-lapse phase contrast images of E. coli cells in a single trap of a mother machine type microfluidic device, (**a, bottom**) Output of cell segmentation of the time-lapse images in (a, top). The cell at the bottom of the kymograph (and thus at the bottom of the cell trap) corresponds to the green trajectory in (b). (**b**) Cell area over time, following only the descendant closest to the bottom, for three examples (in different colors) of cell lineages. (**c**) Growth rates over time for the cell lineages shown in (b). Here the growth rate is calculated for the total cell area of all the descendants in each lineage. Growth rates are estimated using exponential regression of a 10 min sliding window. (**d**) Example of time-lapse phase contrast images and corresponding cell segmentation of S. aureus cells in a single microfluidic trap of size 28x54μm. The cell highlighted in green corresponds to the green curve in (e). (**e-f**) As (b) and (c), but for S. aureus cells. Examples for K. pneumoniae, P. aeruginosa, A. baumannii are shown in Supplementary Figure S1.

## Methods

### Media and batch growth

Overnight cultures (ONCs) were inoculated from glycerol stocks or bacterial plates in Mueller-Hinton (MH) media (70192; Sigma-Aldrich) and grown at 37°C overnight with shaking. For the microfluidic experiments, ONCs were diluted 1:1000 into fresh MH media supplemented with a surfactant (Pluronic F-108; 542342; Sigma-Aldrich) at a final concentration of 0.34 mg/ml and cultured in a 37°C shaker for 2-5 h (duration depended on bacterial strain) before being loaded onto a microfluidic chip. The Pluronic-supplemented MH medium is used throughout all microfluidic experiments and, when indicated, supplemented with antibiotics.

### Microfluidics fabrication

Microfluidics fabrication was performed as previously described in (Baltekin et al. 2017; Kandavalli et al. 2022). Briefly, PDMS [polydimethylsiloxane; Sylgard 184] was cast on a silicon-SU8 mold (ConScience). The silicon wafer has structures of the fluidic channels and modified mother machine traps (trap sizes of 1×1×56μm, 1.125×1.125×56μm, 1.250×1.250×56μm depending on isolate) which were used for the rod-shaped bacteria or wider traps (trap size 1.25×28×54μm) which were used for *S. aureus*. The PDMS chip was punched and bonded to a nr:1.5 glass coverslip (Menzel-Gläser) using plasma treatment (HPT-200, Henniker plasma) followed by incubation at 80°C overnight.

### Cell loading and imaging

Tubings (Tygon) were connected to the punched microfluidic chip using metal tubing connectors. Pressures in the chip were regulated using an OB1-Mk3 (Elveflow). Bacterial cells were loaded onto the chip with growth media flowing through the device at 200 mbar. The microscope enclosure was kept at 37°C. Cells were imaged every minute in phase contrast using custom scripts in Micro-Manager (Edelstein et al. 2010).

### Switching growth media in the microfluidic device

To switch the media that goes into the microfluidic chip, a 3-way split (IDEX) was inserted on the tubing close to the chip media inlet. The third outlet adds a valve-controlled route to a waste container. Switching of growth media in the microfluidic chip was carried out using the following steps: (*i*) The medium in the growth media container was exchanged; (*ii*) The valve on the waste side of the 3-way split (IDEX) was opened, which allowed the old media to be flushed from the tubing directly to waste; (*iii*) The valve was closed and the new media flowed the last two centimeters from the 3-way split to the cells on the chip. To pinpoint when the new media reaches the cells, a fluorescent probe (Cy3) was added to the media. A subset of cells not used for downstream analysis were imaged in epifluorescence to map the time point when the new media reaches the cells. If two media switches are done in the experiment, the second medium does not contain any fluorescent probe. Here, instead, the decrease in fluorescence was used to indicate when the second medium reached the cells.

### MIC assay

Minimum inhibitory concentration (MIC) assays were performed in flat bottom microtiter plates using broth microdilution (Kadeřábková et al. 2024). Bacteria were grown on LB agar plates or in MH. Bacterial suspension from colonies or logarithmic phase culture were diluted in MH to achieve a final inoculum of 1×10^6^ CFU/mL. Two-fold serial dilutions of antibiotics were prepared in MHB to obtain a range of concentrations by transferring 50 μl of solution from wells in the left to right direction. 50 μl of diluted bacterial solution was added to each well. Plates were incubated at 37°C for 16-20 hours and OD600, used to quantify bacterial growth, was measured using a microplate reader (Tecan Sunrise).

### Bacterial strains

Information about strains used in the study is found in Table S1. Bacterial species of the isolates were confirmed using Bruker MALDI Biotyper at the Akademiska hospital in Uppsala. Bacterial species of the isolates were confirmed using Bruker MALDI Biotyper at the Akademiska hospital in Uppsala.

### Microscope setups

Phase contrast and epifluorescence images were acquired using either a Ti2 (Nikon) or a Ti-E (Nikon) microscope. The Ti2 microscope was fitted with a CFI Plan Apo lambda 1.45/100x oil DM (Nikon) objective, a DMK 38UX304 (The Imaging Source) camera, and a

Spectra Gen. 3 (Lumencor) light source for epifluorescence. For Cy3 epifluorescence, a filter cube with FF562-Di03 (Semrock) dichroic mirror, FF01-543/22 (Semrock) excitation filter and FF01-586/20 (Semrock) emission filter was used. The Ti-E microscope was fitted with a CFI Plan Apo Lambda 1.45/100x oil (Nikon) objective, a Ph3 phase plate (Nikon), a DMK 38UX304 (The Imaging Source) camera, and a Sola FISH (Lumencor) light source for epifluorescence. For Cy3 epifluorescence, a filter cube with F555-Di03 (Semrock) dichroic mirror, FF01-530/11 (Semrock) excitation filter and FF01-575/19 emission filter was used. Both microscopes were enclosed in a Lexan incubator (OKOlab) where the temperature is kept constant.

### Image analysis

The acquired phase contrast images were segmented using an Omnipose deep neural network (Cutler et al. 2022), which has been trained using outlines generated from fluorescently labelled *E. coli, A. baumannii* and *S. aureus* cells in mother-machine traps and manually labelled outlines of mixed species data from (Kandavalli et al. 2022) and (Cutler et al. 2022). The fluorescent cells were segmented using either the LoG filter or ilastik (Berg et al. 2019). The training dataset consisted of 1100 images containing 162222 cells. Segmented cells grown in the mother machine type microfluidic device (*K. pneumoniae, K. variicola, P. aeruginosa, A. baumannii, E. coli*) were tracked from frame-to-frame using the Baxter algorithm (Magnusson et al. 2015). *S. aureus* cells grown in a 28x54μm microfluidic trap were tracked using TRACKASTRA (Gallusser and Weigert 2025). A pretrained Omnipose model bact_phase_affinity was used as a starting point to train a model for *S. aureus* cells on a dataset of manually labelled *S. aureus* cells (839 cells in 35 images) and mixed species data from (Cutler et al. 2022) resulting in a total of 284 images with 28357 cells used for training.

### Post-processing of segmented and tracked cells

#### Lineage validity

Within each growth channel, candidate mother cells are identified as those at the closed end of the channel. A channel is retained as a valid lineage only if at least one of its identified mother cells is of good enough segmentation quality and has a pre-switch growth rate of at least 0.002 min^-1^. The earliest such mother cell also needs to be born 10 min before the start of the antibiotic treatment. For the cell size estimations used together with antibiotics that affect cell size and/or lysis, it is required that the mother cell has divided before the switch so that a pre-switch baseline size is available. Valid lineages are then passed to the growth rate and cell size estimation steps described below.

#### Growth rate estimates

Growth rates were estimated using exponential regression within a sliding window of up to 10 minutes, when 10 minutes of preceding data was available. The sliding window regression was applied to trajectories over entire cell lineages where, after cell division, the areas of the descendant were summed. When a cell lineage did grow out of the trap, the lineage was restarted from a descendant of the mother cell closest to the trap constriction. For *S. aureus* the cell area trajectories have large shifts in the cell area at division. These shifts in size at division are an artifact of the fact that the phase contrast images are a 2D projection of a 3D volume, since it has been shown that the volume of the mother cell and the summed volume of the two daughter cells does not change upon division (Zhou et al. 2015). To remedy this size bias at division growth curves of *S. aureus* were smoothed by multiplying all post division sizes by a constant such that the ratio of the size at the last frame before division and the sum of the areas of the descendants after division is the same as the average frame-to-frame size ratios of four frames centered around the division time. Mean growth rates were measured for each cell during 50 minutes prior to switching to antibiotic-containing medium. To remove non-growing segmented objects, traps in which the mean growth rate was less than 0.002 min^-1^ were excluded from further analysis. Cell-specific growth rates were normalized to the mean growth rate measured in the corresponding cell lineage.

#### Cell size estimates

The relative cell area trajectories in Figures 3, 5 and 6 were calculated as follows. Within each valid growth channel, a baseline cell is identified as follows: if no division occurs within half a generation time after the media switch, the last cell at the bottom of the microfluidic trap to divide before the media switch (in sequential experiments, the switch to the second antibiotic) is set as the baseline cell. If, instead, a division occurs within half a generation time after the media switch, the baseline cell is set to the cell at the bottom of the trap before this division. The frame of division for the baseline cell was taken as the alignment point t = 0. The area of the baseline cell at its final frame was used as the baseline area for normalisation. A single-cell growth rate was estimated for the baseline cell by fitting a straight line to the natural logarithm of its segmented area over time and the generation time was computed as ln(2) divided by this rate. Channels for which a generation time could not be estimated were excluded.

The full area trajectory for each trap was assembled from the tracked cells by joining the measured areas of the mother cells and of their tracked descendants. At each division, the daughter remaining at the closed end of the channel was the descendant being followed. When the tracked trajectory ended before the end of imaging, but at least 5 minutes after the switch, it was extended to the end of the experiment by estimating a pseudo cell area as the channel trap area divided by the smoothed cell count in that channel or as the channel trap area when the channel was empty.

Each area was divided by the baseline area to give the relative cell area. Time was normalised to generations by dividing the time relative to t = 0 by the generation time estimated from the last dividing mother cell before the media switch.

### PCA and susceptibility classification

#### Trajectory representation

Each cell lineage was represented as a trajectory vector. For growth rate analyses, the vector was the relative growth rate at regular timepoints (minutes) aligned so that antibiotic addition defined t = 0. For cell area analyses, the vector was the natural log of relative cell area sampled on a generation normalized time, tau, binned at 1/15 resolution and aligned as described above to define tau = 0. In both cases, trajectories were assembled into a lineage x timepoint matrix and restricted to a defined analysis window.

#### Lineage filtering

Within each analysis window, only lineages with complete (100%) coverage of the window’s timepoints were retained so that every lineage contributed a complete feature vector. For area analyses, it was tracked whether any bin within the window had a pseudo-size included.

#### Dimensionality reduction

Principal component analysis (PCA) was performed on the training lineages only and applied unchanged to the test lineages. Trajectories were mean centered using the training mean. PCA was performed on the centered data, so the components reflect the dominant directions of variation in the trajectories.

#### Logistic regression

A logistic regression was trained on the first two principal components with class weight set to “balanced” so that each class contributed equally to the loss regardless of the resistant/susceptible ratio in the training set. Lineages were assigned to a class at the probability threshold of 0.5. Resistant was the positive class in every analysis.

#### Validation schemes

The same classifier was evaluated under the three validation schemes described below.

#### Held-out replicate (Figure 2 and 3)

**Figure 2.**
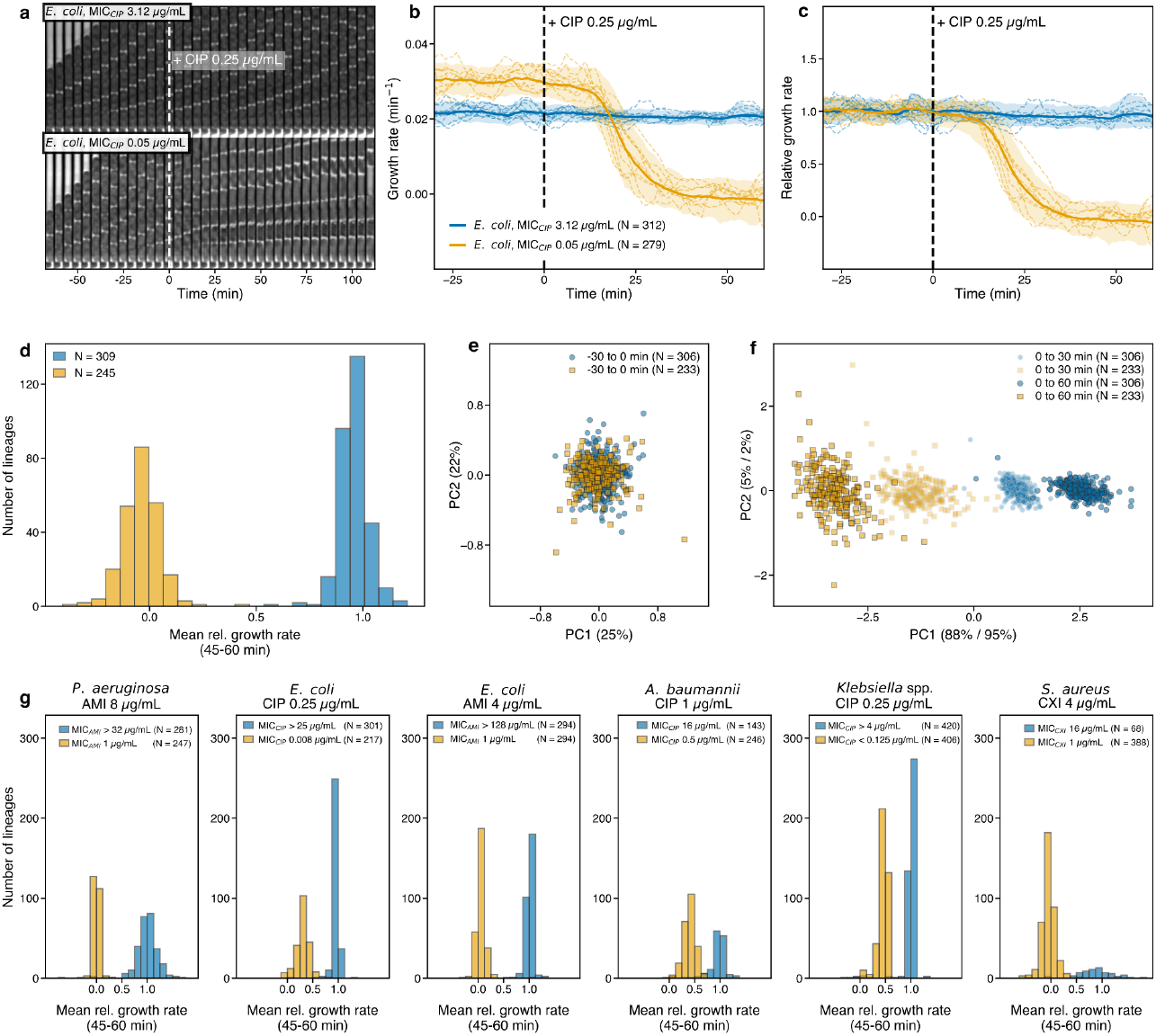
Susceptibility classification of cells after treatment with growth rate affecting antibiotics. (**a**) Kymographs showing time-lapse phase contrast images for two example lineages from a resistant (MIC_CIP_ 3.12 μg/mL) and susceptible (MIC_CIP_ 0.05 μg/mL) isolate of E. coli respectively. CIP at 0.25 μg/mL is added to the medium at dashed white line. (**b**) Growth rate before and after addition of 0.25 μg/mL CIP to the growth medium for resistant (blue) and susceptible (orange) isolates of E. coli. Tinted regions are within limits set by 5% and 95% quantiles of the growth rate distribution for each time point. Solid lines are average growth rates. Dashed lines are 10 randomly selected example lineages. (**c**) Growth rates after normalizing each trajectory in (b) to its pre-treatment baseline. (**d**) Distribution of normalized growth in (c) rates averaged over timepoints from 45-60 minutes. (**e**) Visualization of the first two principal components of a PCA of growth trajectories in (c) before antibiotic treatment. (**f**) Visualization of PCA at two different time intervals after antibiotic treatment shown in (c). (**g**) As (d) but for different strains at different antibiotics or antibiotic concentrations as indicated in the figure. Klebsiella spp. indicates a combination of K. pneumonia (resistant) and K. variicola (susceptible).

**Figure 3.**
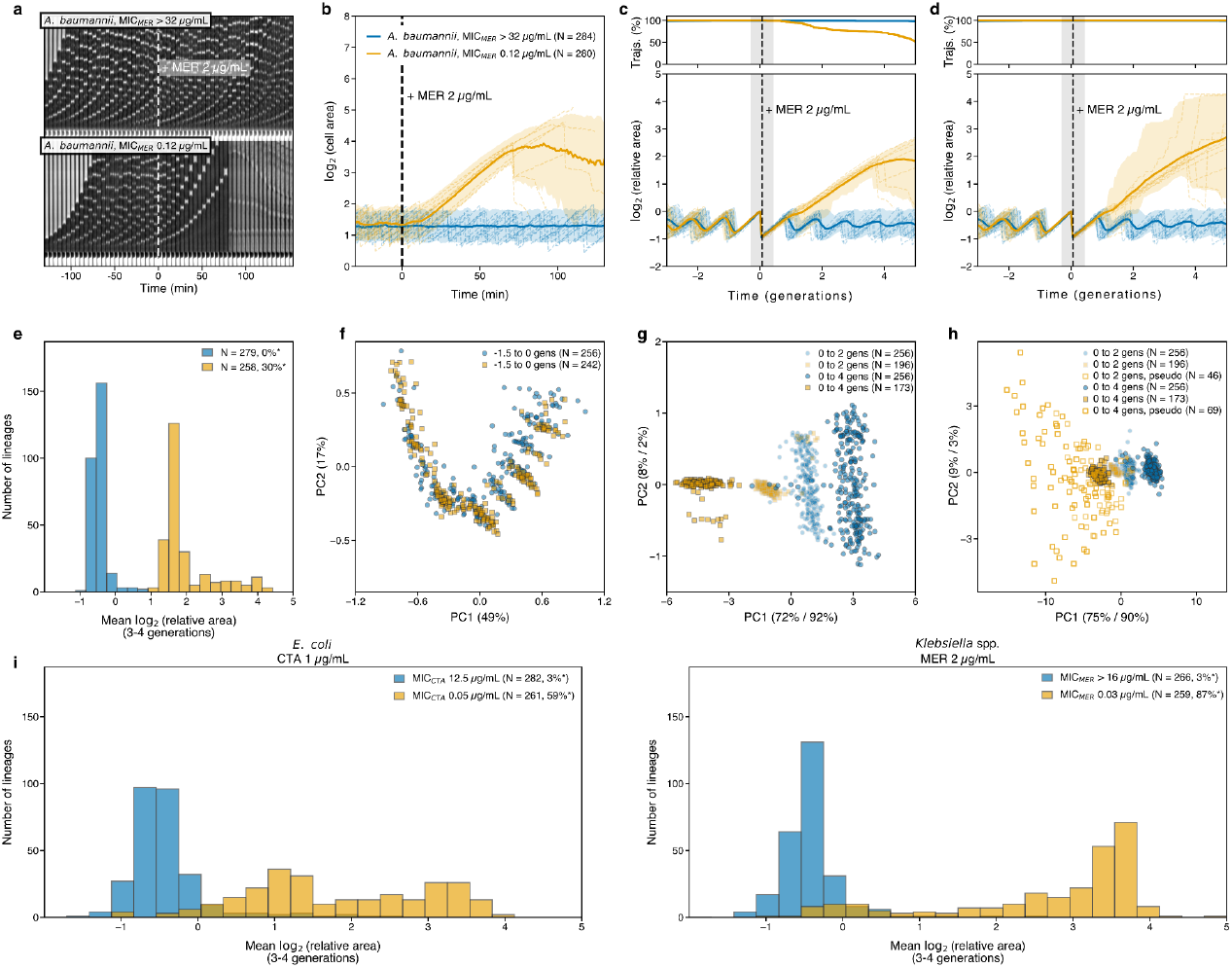
Susceptibility classification of cells after treatment with antibiotics causing cell size increase or lysis. (**a**) As Fig. 2a, but for A. baumannii treated with MER. (**b**) Growth and division for single lineages where at divisions one of the descendants are tracked before and after treating A. baumannii with MER. Coloring and markers as in Fig. 2b (**c, bottom**) Data in (b) normalized to their corresponding pre-treatment baseline. Here the area is relative to the last division area before MER treatment, time is relative to the time of last division before MER treatment and the scale of the time-axis in the unit of pre-treatment generation time for each lineage. Colors and markers as in Fig. 2c, but here the dashed vertical black line shows the average time of antibiotic treatment and the gray shaded area show one standard deviation of the average time distribution in antibiotic treatment time. (**c, top**) Fraction of growth trajectories still remaining in the analysis. The main reason for trajectory loss is due to cell lysis. (**d**) Data in (c) in which lysing cells have been assigned a pseudo-size, based on the fraction of lysed cells in each cell trap. See main text and methods for detailed definition. Example trajectories with pseudo sizes after lysis are shown with dash-dotted curves. (**e**) Distributions of the mean relative cell size between 3 and 4 pre-treatment generation times after antibiotic treatment for cells from susceptible and resistant isolates show in (d). (**f**) As Fig 2e, but for data shown in (c). (**g**) As Fig. 2f, but for data shown in (c) (**h**) As Fig. 2f, but for data shown in (d). (**i**) As (d), but for E. coli treated with CTA and K. pneumoniae treated with MER.

Within a group of biological replicates of one species/antibiotic condition, the classifier was trained on all but one replicate and tested on the held-out replicate. The replicate shown in the main figures was always retained in the training set and never used as the test replicate. The test replicate was drawn at random from the remaining replicates. PCA was fit only on the pooled training replicates.

#### Pooled cross-validation (Figure 5)

Lineages from all experiments of a given condition were pooled and evaluated by 5-fold cross-validation with PCA refit within each training fold. Reported AUC, accuracy, sensitivity and specificity are computed on the out-of-fold predictions.

#### Train on monotherapy, test on sequential (Figure 6)

In the sequential experiments, each stage was classified separately. For each stage, a classifier was trained on independent monotherapy experiments using only that antibiotic and then used to predict resistance/susceptibility at the corresponding stage of the sequential experiment. So the sequential data was only used as a test set.

## Results

### Establishing the phenotypic baselines for cells before antibiotic treatment

The first question is how long should be spent estimating the untreated phenotype to determine when it can be used as a stable baseline for subsequent antibiotic treatment. We will initially describe the case of antibiotics that impact growth rate, such as fluoroquinolones and aminoglycosides. Growth-rate trajectories of three examples from an isolate of *E. coli*, and three examples from an isolate of *S. aureus* are shown in Figures 1c and 1f, respectively. Here a cell and its descendants, *i*.*e*. a lineage, is tracked while growing and dividing without antibiotic treatment. For an individual cell, how rapidly the growth rate is randomized during steady state growth can be described by the growth-rate autocorrelation. The decay in autocorrelation for isolates of *K. pneumoniae, P. aeruginosa, A. baumannii, E. coli and S. aureus* are shown in Supplementary Figure S2. In summary, the autocorrelation decays rapidly within the first 10 minutes, which we attribute to short time scale noise in the growth-rate estimates. Note that the rate of this decay is dependent on the size of the time window used for growth rate estimates, which is set to 10 minutes (Methods). At approximately 1 generation time, most strains show a peak in the auto correlation, which is especially prominent in *S. aureus*. The round shaped *S. aureus* previously has been shown to grow exponentially in volume (Zhou et al. 2015) in between division and it is possible the cell cycle dependent variations in our growth rate estimates are due to various biases introduced by the fact that we are using standard phase contrast microscopy to observe a 2D projected area of a true 3D volume. We use 50 minutes of pre-antibiotic growth to average out both the short time scale fluctuations and cell cycle dependent biases. This time also allows for checking that the cells are growing at steady-state before antibiotic is added to the growth medium.

For antibiotics that affect the cell size, the pretreatment baseline is the cell division size. In line with previous observations (Deforet et al. 2015; Taheri-Araghi et al. 2015; Campos et al. 2014), we find that autocorrelation of division sizes decreases slowly over multiple generations. The time scale for this decrease is longer than the expected phenotypic response time of antibiotic treatment. Based on these observations, the last cell division before antibiotic treatment is more informative for the expected size in the next division than the average division size. For this reason the last division size before antibiotic treatment is used as the baseline phenotype for cell size affecting antibiotics and thus the cell has to be observed for the time of one generation in order to ensure that it has divided at least once. The phenotype for antibiotics that cause rapid lysis is also linked to cell size, since the same antibiotic, for example, Meropenem, can cause either cell size increase or rapid lysis depending on the species it is applied to and the antibiotic concentration used (Kim et al. 2023). For this reason lysis and cell size increase is analyzed together. The lysis frequency in absence of antibiotics is expected to be very low, however, imperfections in the cell tracking analysis may lead to loss of cell trajectories which are incorrectly interpreted as cell lysis. The loss frequency during 30 minutes prior to antibiotic treatment was found to be 1.2% on average, but varies between different isolates (Supplementary Table S2). Given that the cell needs to divide at least once to establish the division size baseline and that the lysis analysis improves with more descendants in the trap, the time it needs to be observed before treatment is in the order of two generations.

### When can an individual cell be used to classify an isolate as susceptible?

Having established the phenotypic pre-treatment baseline, the next question is if it is possible to conclusively classify an isolate as susceptible based on the information from a single cell, and if so, how much time is required to do so. We first exemplify the principles of the classification by using isolates with a MIC that are well separated from the antibiotic concentration used in the test. The antibiotic test concentrations are typically set to the EUCAST breakpoint for each combination of species and antibiotic. For the two other cases (*E. coli* with AMI & *P. aeruginosa* with AMI) the antibiotic test concentrations are set one log2 step below the breakpoint, which does not influence the interpretations in these cases since the MICs of the strains are well separated from the antibiotic test concentrations. Isolates with MIC values above and below the breakpoints are referred to as resistant and susceptible, respectively.

To obtain a first-order estimate of how well the susceptibility classification performs, we tested highly resistant and susceptible isolates of *E. coli, P. aeruginosa, S. aureus, A. baumannii* and a combination of *K. pneumonia* (resistant) and *K. variicola* (susceptible), using antibiotics that affect growth rates, cell size, and/or cause cell lysis. The combination of *K. pneumonia* and *K. variicola* will be referred to as *Klebsiella spp*. The number of drug-bug combinations that one could potentially test rapidly becomes intractable, instead our selection was made to illustrate a diverse set of examples which are clinically relevant for BSI.

### Susceptibility classification for antibiotics affecting growth rates

Figure 2a-b shows the growth rate responses for cells from two different isolates of *E. coli* treated with ciprofloxacin (CIP). Here growth rate traces from the resistant isolate are shown in blue and traces from the susceptible isolate are shown in orange. Prior to antibiotic treatment, cells from the resistant isolate grow slower as compared to cells from the susceptible isolate. After antibiotic treatment the cells from the susceptible isolate decrease in growth rate while cells from the resistant isolate remain approximately constant (Figure 2b). The results illustrate the importance of comparing the growth rate response to the pre-treatment phenotypic baseline as different isolates can have very different pre-treatment growth rates and Figure 2c shows the growth rate response to CIP normalized to its pre-treatment baseline. Using principal component analysis (PCA) we show that the individual resistant and susceptible cells form distinct clusters in the two first principal vectors due to antibiotic treatment (Figure 2e-f) and that the distance between the clusters increases with increasing time after antibiotic treatment (Figure 2f). Using logistic regression on the clusters in the first two principal coordinates, we construct a classifier for calling an isolate as being susceptible or resistant. We evaluate the performance of the classifier at different durations of antibiotic treatment in a repeat experiment using the area under the curve (AUC) of a true resistant vs false resistant plot. We find that the AUC increases to close to 1 after 30 minutes after which it stays constant over time (Supplementary Figure S3). For example, after 45 minutes of antibiotic treatment the risk of incorrectly classifying a single cell lineage from a resistant isolate as being susceptible can be kept to 0.3%, while at the same time the risk of incorrectly classifying a single cell lineage from a susceptible isolate as resistant is 0.7% (Supplementary Table S3). We performed the same type of experiments on *E. coli, P. aeruginosa, Klebsiella spp*., *S. aureus, and A. baumannii* treated with CIP, Amikacin (AMI), or Cefoxitin (CXI), where in each case the single cell lineages distributions of time-average growth rates 45-60 minutes after antibiotic treatment separate well between resistant and susceptible isolates (Figure 2g). For the drug-bug combinations shown in Figure 2g, the risk of misclassifying a single-cell lineage from a resistant isolate as susceptible is at most 1.8%, while the risk of misclassifying a single-cell lineage from a susceptible isolate as resistant is at most 8.3%, within 35 to 60 minutes of treatment depending on species (Supplementary Table S3).

### Susceptibility classification for antibiotics causing cell size increase and lysis

In addition to experiments and analysis with growth rate affecting antibiotics, we performed corresponding experiments and analysis with antibiotics that inhibit cell division and/or cause cell lysis. Here we track the cells at the bottom of each cell trap and observe how they grow and divide over time, before and after treatment with antibiotics. Figure 3a, b shows growth and division for *A. baumannii* cells from one resistant and one susceptible isolate treated with meropenem (MER). Growth and division traces for cells from the resistant isolate are shown in blue and traces from the sensitive isolate are shown in orange. Here the cells from the susceptible isolate keep growing past their expected division size, while the cells from the resistant isolate keep growing and dividing in the same manner as before antibiotic treatment (Figure 3b). To evaluate if cells are deviating from their untreated division pattern, the cells were synchronized in time and normalized in size to the last division event before antibiotic treatment (Figure 3c). By PCA analysis we show that the growth and division trajectories from the susceptible and resistant isolates form distinct clusters already before antibiotic treatment (Supplementary Figure S4A). The observed clustering is due to the growth rate difference of the two isolates. By normalizing the time scale of each growth and division trajectory to its own pre-treatment generation time (Figure 3g & Supplementary Figure 4B), the clustering after PCA is solely based on response to antibiotics (Figure 3f-g). In addition to cell filamentation in response to MER, some of the *A. baumannii* cells also lyse. The lysis phenotype is even more pronounced in Klebsiella spp., where a large fraction of the cells lyse before any change in cell size can be observed. Since cell filamentation and lysis appear together, our analysis and classification account for both phenomena jointly in the following manner. If a cell is lost in cell tracking, presumably due to lysis, its observed and recorded cell size is replaced by a pseudo-size. The pseudo cell-size is defined as the product of the total microfluidics trap size and the fraction of the total number of cells that remains in the trap as compared to when the track was lost, *i*.*e*. effectively interpreting lysis as an extreme filamentation. The result of the combined analysis is shown in Figure 3d.

As in the growth-rate inhibiting antibiotics above we construct a classifier based on logistic regression and apply this to repeat experiments to evaluate its performance. We find that for the AUC to reach close to 1 requires a time corresponding to 2.5 pre-treatment generations of MER treatment for the *A. baumannii* isolates in Fig. 3h. Here, for example, after antibiotic treatment for 3 pre-treatment generation times, 1.8% of the single cell lineages from the resistant isolate cells were misclassified as susceptible, while at the same time 3.2% of the single cell lineages from the susceptible isolate were misclassified as resistant. Additionally we performed the same type of experiments for isolates of Klebsiella spp. treated with MER and for isolates of *E. coli* treated with cefotaxime (CTA) where we find that the cell size distributions separate well 3-4 generation times after antibiotic treatment (Figure 3i). We performed the same classification methods on these two combinations of susceptible and resistant isolates. Here we found that for Klebsiella spp. treated with MER the risk of incorrectly classifying a single lineage from the resistant isolate as sensitive is 3.2% after antibiotic treatment duration corresponding to 4 pre-treatment generations while at the same time the risk of incorrectly classifying a single cell lineage from the sensitive isolate as resistant was 7.2%. In the *E. coli* isolate which is resistant to CTA, there is a subpopulation of cells that respond to CTA treatment in the same manner as cells from the CTA sensitive isolate. This leads to the high risk (12.4%) of incorrectly classifying a cell lineage from the resistant isolate as sensitive after 3 pre-treatment generations. The impact of the heterogeneity on the single cell pAST is described further in the discussion.

### Operating close to the breakpoint

In the examples provided so far, the cells from the resistant isolates have largely been unaffected by the antibiotic treatment and thus it may seem sufficient to observe a response that is statistically significantly different from the pre-antibiotic treatment in order to classify an isolate as susceptible. However, some resistant isolates, especially those with MICs near the antibiotic concentrations used in the test, may respond to antibiotic treatment to some extent. This means that, for each antibiotic testing concentration used, a classifier needs to be trained on sufficiently many isolates of different MICs in order to capture the full range of possible single-cells responses.

To exemplify the single cell responses for cells from isolates of varying MIC treated with the same antibiotic concentration, we grew four different isolates of *E. coli* with MICs 2, 8, 32 and >128μg/ml treated with 8μg/mL of amikacin (AMI), which is the EUCAST breakpoint for clinically susceptible. Thus two of the isolates are resistant and two are susceptible. Figure 4 shows the distributions of single cell growth rates at different time points after antibiotic treatment. The isolates with MICs of 2μg/mL and 128μg/mL, being either well below or well above the testing concentration, respectively, behave as the previous examples in Figure 2. The isolate with MIC 8μg/mL responds more slowly to the antibiotic treatment, but has after 90 minutes a very similar distribution of growth-rates as the isolate with MIC of 2μg/mL. For the MIC 32μg/mL isolate there is heterogeneity in the response. The majority of the cell lineages respond in a similar manner as the cells from the 8μg/mL isolate, while a subpopulation of cell lineages retain a growth-rate similar to the untreated case.

**Figure 4.**
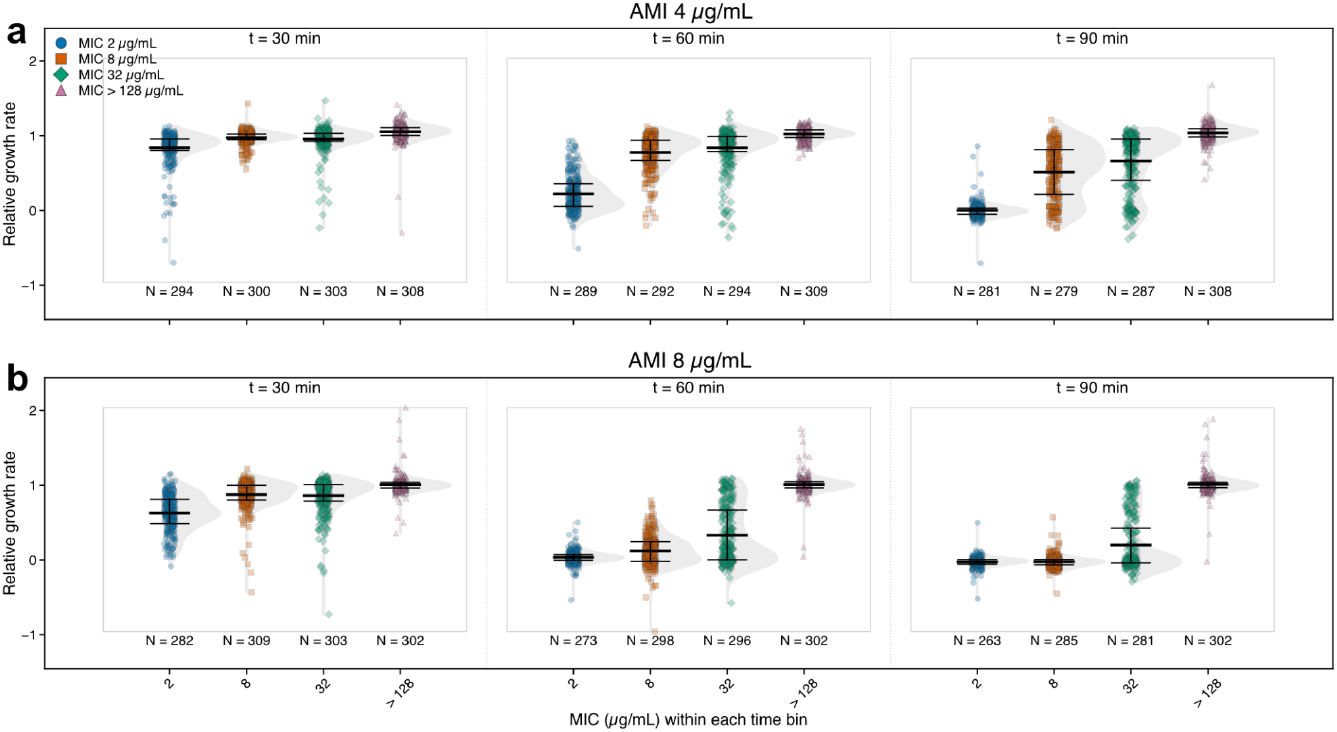
Relative growth rates across AMI MICs and treatment times. Raincloud plots visualizing distributions of relative growth rates of four E. coli isolates with different MICs (as indicated in the figure) measured 30, 60 and 90 min after AMI introduction. The concentration of AMI used was 4 μg/mL in (a) and 8 μg/mL in (b). Grey outlined boxes group the four isolates at each post-treatment time point. Relative growth rates were calculated as in Fig. 2 by normalizing each lineage’s growth rate to its time-averaged pre-treatment growth rate. Each plot combines a half-violin representing the distribution, individual data points and a boxplot summarizing the mean and interquartile range (25th-75th percentiles).

This raises an important question: why do a majority of the cell lineages stop growing when treated at concentrations below the isolate’s MIC? A likely explanation is the obscurement of the single-cell perspective in the conventional bulk MIC assay. In bulk, it is sufficient for a small subpopulation of cells to continue growing for the culture to be scored with a high MIC. As a result, the assay reports the survival of the few rather than the fate of the many, whereas single-cell measurements reveal the true heterogeneity and time dependence of the antibiotic response. Having the possibility of characterizing the distribution of single cell behaviours in the isolate, is, however, not helpful in the case of BSIs where there is potentially only a single cell to analyse. If this single cell stops growing despite being sampled from an overall resistant isolate, it would be misclassified. For this reason, it may be needed to calibrate the single-cell assays to work with sub-breakpoint concentrations, at the cost of misclassifying some susceptible as resistant. We exemplify this by treating the cells from the four different isolates with 4μg/mL of AMI instead of 8μg/mL (Figure 5b). Except for the isolate with the highest MIC, the response to 4μg/mL of AMI is decreased at all timepoints as compared to the 8μg/ml case. At 60 minutes after antibiotic treatment a substantially smaller fraction of the cell lineages from the 32μg/mL isolate have a very slow growth rate, and would be misclassified as resistant. This is also true for the susceptible 8μg/mL isolate, however, since its MIC is at the breakpoint this is as expected.

**Figure 5:**
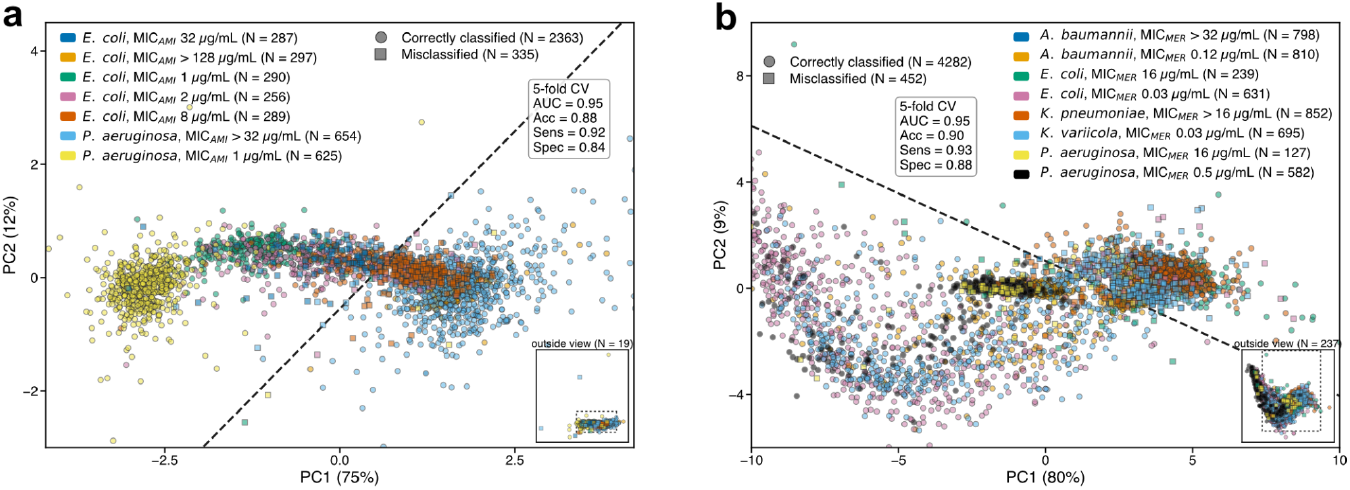
(**a**) Visualization of the two principal components of a PCA of the response to 45 minutes of AMI (8μg/mL) treatment together with decision boundary (dashed black line) from a logistic regression based classifier on seven different strains as indicated in the figure with different colors. Correct classification is indicated by a circle and incorrect classification is indicated by a square. Limits on PC1 and PC2 axes have been truncated to highlight details of the cluster. The full data set is shown in the inset in the lower left corner. Sensitivity and specificity are defined as the true resistant and the true susceptible rates, respectively. (**b**) As in (a), but for MER (2μg/mL) treatment during a time corresponding to 3 pre-treatment generations.

### Working with multiple species and multiple MICs and a single antibiotic concentration

Having characterized the behaviour of isolates with MICs close to the testing concentration, we aimed to benchmark the classifier at these more difficult cases. A practically functional susceptibility classifier also needs to handle different species, since different strains may have different breakpoints for the same antibiotic. If the species can be identified to some degree before the antibiotic exposure, the problem is greatly simplified for antibiotics where the breakpoint is similar for phenotypically similar species (https://www.eucast.org/fileadmin/eucast/pdf/breakpoints/v_16.0_Breakpoint_Tables.pdf). To exemplify a case in which the most important BSI-related rod shaped species share a similar breakpoint (8 μg/mL for *E. coli, A. baumannii and K. pneumoniae* and 16 μg/mL for *P. aeruginosa)* for AMI, we grew two isolates *P. aeruginosa* with AMI MICs >32μg/mL and 1μg/mL and treated these with 8μg/mL of AMI. The data from these experiments were combined with data from the four *E. coli* strains presented above and an *E. coli* isolate of AMI MICs 1 μg/mL. A classifier was constructed in the same manner as in Figure 2 based on PCA and logistic regression (Figure 5a). We evaluated the performance of the classifier by doing 5 rounds of training on 80% of the data and evaluating on the remaining 20%, where each round generates a different 80/20 split. Here we find that the AUC=0.95 after 45 minutes of antibiotic treatment, and at this time point the risk of misclassifying a single cell lineage from either of the resistant isolates as susceptible is 8% while at the same point the risk of incorrectly classifying a single cell lineages from either of the susceptible isolates as resistant is 16% (Supplementary Table S4). As expected the majority of the misclassified cell lineages stems from the two *E. coli* isolates with MIC 8μg/mL and 32μg/mL. For the MIC 8μg/mL isolate which is on the breakpoint, the single cell population gets split into ∼30% susceptible and ∼70% resistant. More unexpectedly, the MIC 32μg/mL isolate also gets split into ∼30% susceptible and ∼70% resistant. The unexpectedly large fraction of single cells being classified as susceptible stems from the fact that, on the single cell level, these cells behave similar to cells from the 8μg/mL isolate, which is indeed susceptible (Supplementary Figure S5). It should also be noted that isolates with MIC values near the clinical resistance breakpoint are often difficult to classify, also using millions of cells, since even the gold-standard MIC assay has an uncertainty of approximately one two-fold dilution (±1 log_2_ unit) (Kahlmeter and Turnidge 2023).

We also performed susceptibility classification for a joint data-set with resistant and susceptible isolates from *A. baumannii, E. coli, K. pneumoniae* and *P. aeruginosa* which all share the same breakpoint and treated these with 2ug/ml of MER (Figure 5b) which induces cell size increase and cell lysis. Here we find that after antibiotic treatment time corresponding to 3 pre-treatment generation times, the risk of misclassifying a single cell lineages from the resistant isolate as sensitive is 7%, while at the same time the risk of misclassifying a single cell lineages from the susceptible strain as resistant is 12%.

### Sequential antibiotic treatments does not impact susceptibility classification efficiency

Having classified a single cell lineage as being resistant, it is however still unclear which antibiotic to use for a patient. Thus, the antibiotic, or the antibiotic concentration will need to be changed and a test carried out with this new antibiotic or antibiotic concentration. A caveat of this approach is that a tested cell’s response to an antibiotic may depend on prior treatments of the same cell lineage. To exemplify how the sequential treatments could work, we used two *E. coli* isolates where one is resistant to CIP and susceptible to AMI and the second isolate is resistant to both AMI and CIP. The susceptibility classification of the two isolates to only AMI treatment was already tested in Fig 2. In Figure 6a-c we show that the cell’s response to AMI is not affected if we pre-treat with CIP. The susceptibility classification performance is still very high where the sensitivity and specificity stays approximately the same (Supplementary Table S5). Figures 6d-f show the result of the corresponding experiments using CTA. Here, the sensitivity and specificity of the susceptibility classification changes from 0.88 and 0.98 in the single treatment case (Supplementary Table S2) to 0.90 and 0.93 in the sequential double treatment case (Supplementary Table S5). Thus, for the two examples presented, antibiotic treatment can be applied sequentially without any substantial loss in the performance of the susceptility classification.

**Figure 6.**
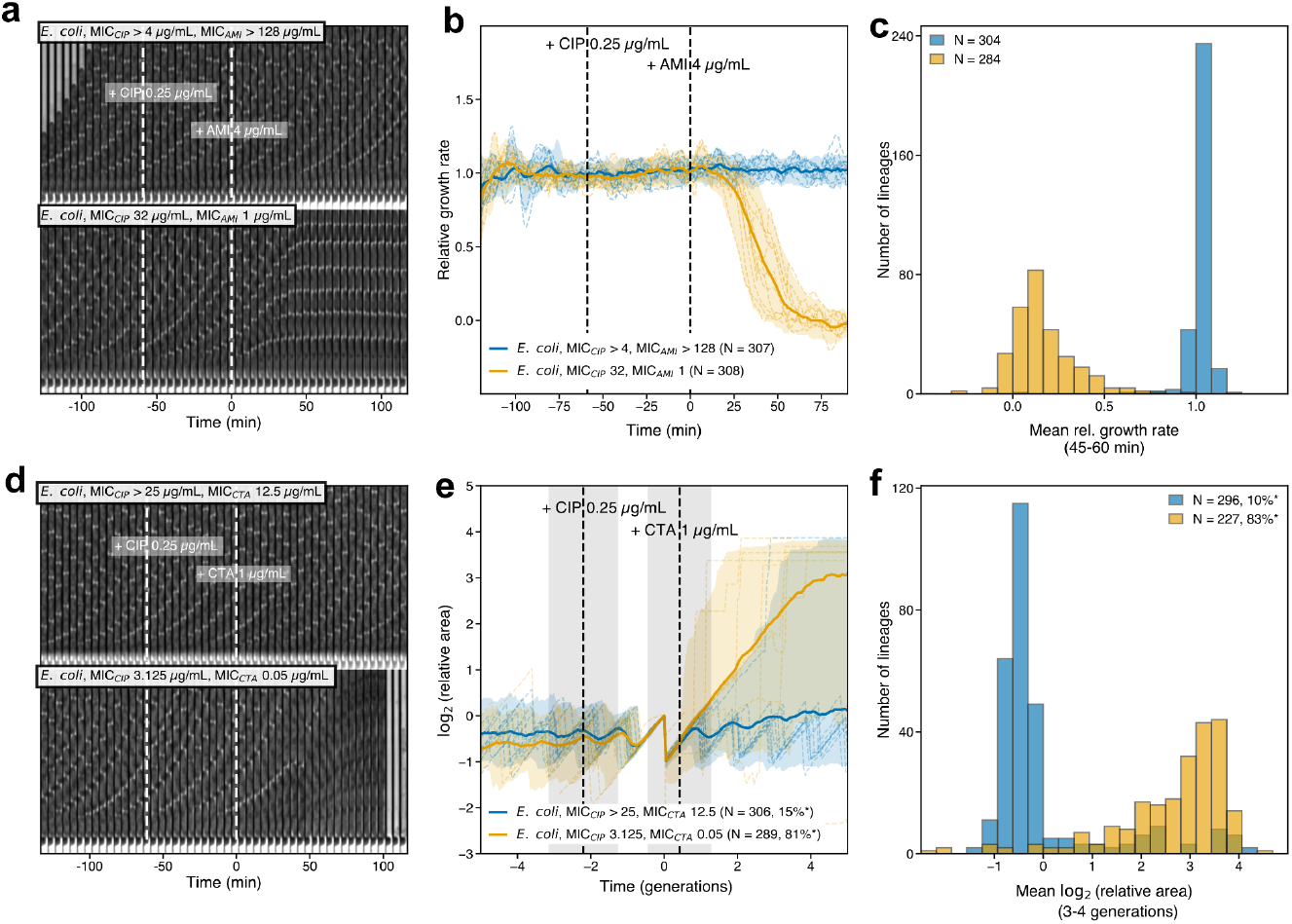
The effect of sequential antibiotic treatment. (**a**) As Fig 2a, but with CIP (0.25μg/mL) and AMI (4μg/mL) treatment of two strains being either resistant (MIC as indicated in figure)) or susceptible to AMI. The two strains are the same as the AMI treated E. coli shown in Fig 2. (**b**) As Fig. 2c, but with sequential CIP and AMI treatment of the strains in (A). (**c**) As Fig 2d, after 45-60min of AMI treatment. (**d**) As (a), but with CIP (0.25ug/ml) and CTA (1ug/ml) treatment of two strains being either resistant or susceptible to CTA. The two strains are the same as the CTA treated E. coli shown in Fig 3. (**e**) As Fig. 3d, but with sequential CIP and CTA treatment of the strains in (d). (**f**) As Fig 3e, after CTA treatment for a time corresponding to 3-4 pre-treatment generation times.

## Discussion

We have outlined the possibilities and challenges of performing pAST based on data from a single bacterial cell. The ideas have been evaluated using the antibiotics ciprofloxacin, cefotaxime, cefoxitin, meropenem and amikacin on rapidly growing *E. coli, P. aeruginosa, K. pneumoniae, K. variicola, S. aureus, and A. baumannii* . For antibiotics relevant to treating *E. coli*, including CIP, we have previously shown that AST can be performed in ≈15 min (Baltekin et al. 2017). However, such assays rely on the average of a few hundred bacteria. For single-cell pAST, on the other hand, additional time is needed to ensure that the distribution of single cell’s responses to antibiotic treatment of the susceptible isolate, including cell-to-cell variability and measurement noise, is different enough from the distribution of responses in the resistant isolates. When this is the case, information from a single cell lineage can be used to assess the susceptibility of the entire isolate. We have shown that for pairwise comparisons of resistant and susceptible isolates of the same species that have MICs which are far from the breakpoint, it is possible to perform accurate single-cell phenotypic AST within 2 hour for growth rate affecting antibiotic (Fig 2) and within 5 cell generations for antibiotics causing cell size increase and/or lysis (Fig. 3). For the cases of antibiotics that affect the growth rate (Fig. 2), *S. aureus* was one of cases with the overall lowest performance. One reason for the lower performance is the more prominent regular temporal fluctuations in growth rate which were observed for *S. aureus* (Fig. 1). The growth of S. aureus has previously been shown to be exponential (Zhou et al. 2015) and we do not believe that the growth rate variability we observe stems from a true variability in growth rate, rather that it is due to systematic, cell cycle dependent, biases in how we determine the cell size. By improving the true 3D size estimate, for example, with quantitative phase contrast microscopy, the performance of the classifier could also be improved.

While classification of resistant and susceptible isolates with MIC far from the breakpoint is a necessity for a functional AST, it must also handle multiple species and isolates with various MICs. As expected, the misclassification rate increases with decreasing distance between the isolate’s MICs and the breakpoint (Fig. 4,5). To avoid misclassifying resistant isolates as susceptible one could shift the decision boundary to be more stringent about calling susceptible cells or lower the test concentration for the antibiotic (Fig. 5). Both are at the cost of more frequently calling susceptible isolates as resistant. Additionally, there is also room for more advanced classification methods, for example based on direct classification of the image sequence of the cell before and after exposing it to antibiotics using AI. Also the duration of the antibiotic treatment could be optimized for maximal classification performance. We also show that single cell pAST works for multiple species that share the similar antibiotic concentration breakpoint (Fig. 5).

This brings us to the challenge that different species have different resistance breakpoints. We have previously shown (Kandavalli et al. 2022) that it is possible to perform a FISH assay to determine species after the AST step, which is a possibility also at the single cell level. Since the FISH assay relies on irreversible fixation of the cells, it is not tractable for choosing the antibiotic test concentration. It can, however, be used for post-test species ID for antibiotics where the cells do not lyse. For selecting the antibiotic test concentration it is possible to use a non-destructive ID method before exposing it to an antibiotic, for example, based on Raman spectroscopy (Ho et al. 2019) or time-lapse phase contrast (Hallström et al. 2023). An alternative is to focus on antibiotics that have the same breakpoint for all relevant bacteria, which we exemplified in the analysis leading up to Figure 5.

For one of the resistant isolates of *E. coli* we find significant cell to cell heterogeneity in the response to CTA. Here a fraction of the cell lineages from the resistant isolate are clearly as susceptible to CTA as cells from the susceptible isolate. Thus, if one of these few cell lineages from the resistant isolates are used for the single cell pAST a very major error will occur. The frequency of this phenomenon has to be characterized in many patients samples, and if it is a general problem for CTA it may imply that this antibiotic is not suitable for a single cell assay unless it’s possible to find adjuvant or growth conditions that reduce the heterogeneity.

The suggested path forward is to work with sequences of drugs that have close to the same breakpoint for morphologically similar bacteria, such as meropenem and tobramycin for rods and linezolid, vancomycin or daptomycin for cocci; and to test many clinical isolates of diverse species with different MIC values such that it is possible to find an optimal classifier based on single cell response trajectories.

In summary, the take-home message is that rapid single-cell phenotypic AST is possible for some, but potentially not all, combinations of drugs and bugs, and that it is important to find which these functional combinations are.

## Acknowledgements

We thank Buu Minh Tran for contributing to initial discussions about the project, Praneeth Karempudi for support with deep learning, Elias Amselem for designing the microfluidic device used for *S. aureus*, and Alice Törnqvist and Kicki Gavalidou at Akademiska hospital Clinical Microbiology laboratory in Uppsala for help with MALDI.

This research was funded by the SSF (grant no. ARC19-0016), the Swedish Research Council (grant no. 2024-06127), the Novo Nordisk Foundation (grant no. NNF23OC0083419), the Knut and Alice Wallenberg Foundation (grant no. 2023.0531), and the eSSENCE e-science initiative. The computations and data management were facilitated by resources from the Swedish National Infrastructure for Computing at UPPMAX, with partial funding from the Swedish Research Council (grant agreement no. 2022-06725). A patent has been filed for the method with a priority date of April 2024.

## Author contributions

IA, LJ & JL made the experiments; SZ & LJ analysed data; JL adapted the fluidic assay; JE, DF & PK wrote the paper with input from the other authors; DF and JE developed statistical reasoning; PK contributed clinical relevance and coordination; JE conceived the idea. Everyone contributed to the evaluation of results, experiments and project planning.

