## Supplementary material for "Phenotypic Antibiotic Susceptibility Testing at the limit of one bacterial cell": SI

### Supplementary Tables:

**Supplementary Table S1. Strain list.** *Data from EUCAST QC Tables v. 16.0 (2026). †MIC value determined by EUCAST. All other MIC values were determined in-house according to the methods section in the Main text.

| **Strain name** | **Species** | **Figure reference** | **Relevant MIC (µg/mL)** | **Strain reference** |
|---|---|---|---|---|
| EL4824 | *E. coli* | Fig. 1, Fig S2, Fig. 2, Fig. S3, Fig. 4, Fig. 5 | AMI 1-2*<br>CIP 0.008* | *Escherichia coli* (Migula) Castellani and Chalmers (ATCC 25922) |
| EL4900 | *E. coli* | Fig. 2, Fig. S3 | CIP 3.12 | Uppsala University Hospital<br>(Baltekin *et al.* 2017) |
| EL4884 | *E. coli* | Fig. 2, Fig. S3 | CIP 0.05 | Uppsala University Hospital<br>(Baltekin *et al.* 2017) |
| EL4668 | *E. coli* | Fig. 2, Fig. 3, Fig. S3, Fig. 6 | CIP > 25<br>CTA 12.5 | *Escherichia coli ALB 53* EUCAST |
| EL4483 | *E. coli* | Fig. 3, Fig. S3, Fig. 6 | CIP 3.12<br>CTA 0.05 | Dan Andersson Lab, Uppsala University |
| EL4987 | *E. coli* | Fig. 2, Fig. S3, Fig. 4, Fig. 5, Fig. 6 | AMI > 128†<br>CIP > 4† | *Escherichia coli*<br>19-597221<br>EUCAST QC strain |
| EL4876 | *E. coli* | Fig. 2, Fig. S3, Fig. 5, Fig. 6 | AMI 1<br>CIP 32 | Uppsala University Hospital<br>(Baltekin *et al.* 2017) |
| EL4871 | *E. coli* | Fig. 4, Fig. 5, Fig. S5 | AMI 32 | Uppsala University Hospital<br>(Baltekin *et al.* 2017) |
| EL4880 | *E. coli* | Fig. 4, Fig. 5, Fig. S5 | AMI 8 | Uppsala University Hospital<br>(Baltekin *et al.* 2017) |
| EL5106 | *E. coli* | Fig. 5 | MER 16 | Helen Wang, Uppsala University |
| EL4988 | *E. coli* | Fig. 5 | MER 0.03† | *Escherichia coli* 8621<br>EUCAST QC strain |
| EL4991 | *K. pneumoniae* | Fig. S1, Fig. S2, Fig. 2, Fig. 3, Fig. S3, Fig. 5 | CIP > 4†<br>MER > 16† | *Klebsiella pneumoniae*<br>TU7<br>EUCAST QC strain |
| EL4990 | *K. variicola* | Fig. 2, Fig. 3, Fig. S3, Fig. 5 | CIP < 0.125†<br>MER 0.03† | *Klebsiella pneumoniae*<br>3560 B |
| EL4649 | *P. aeruginosa* | Fig. S1, Fig. S2, Fig. 2, Fig. S3, Fig. S4, Fig. 5 | AMI 1†<br>MER 0.5† | *Pseudomonas aeruginosa* 75457 from EUCAST collection of defined strains |

| | | | | |
|---|---|---|---|---|
| | | | | https://www.eucast.org/ast_of_bacteria/panels_of_defined_strains/ |
| EL4650 | *P. aeruginosa* | Fig. 2, Fig. S3, Fig. S4, Fig. 5 | AMI > 32†<br>MER 16† | *Pseudomonas aeruginosa* 75458 from EUCAST collection of defined strains https://www.eucast.org/ast_of_bacteria/panels_of_defined_strains/ |
| EL4992 | *A. baumannii* | Fig. S1, Fig. S2, Fig. 3, Fig. S3, Fig. 5 | MER > 32† | *Acinetobacter baumannii* 18-105424 EUCAST QC strain |
| EL4993 | *A. baumannii* | Fig. 3, Fig. S3, Fig. 5 | MER 0.12† | CCUG *Acinetobacter baumannii* AB8 EUCAST QC strain |
| EL4821 | *A. baumannii* | Fig. 2, Fig. S3 | CIP 0.5 | CCUG *Acinetobacater baumannii* 890 |
| EL4670 | *A. baumannii* | Fig. 2, Fig. S3 | CIP 16 | CCUG *Acinetobacater baumannii* CCUG R17 |
| EL3159 | *S. aureus* | Fig. 1, Fig. S2, Fig. 2, Fig. S3 | CXI 1 | *Staphylococcus aureus* ATCC29213 |
| EL4995 | *S. aureus* | Fig. 2, Fig. S3 | CXI > 16 | CCUG *Staphylococcus aureus* 9268 |

**Supplementary Table S2. Loss of single-cell lineage trajectories during analysis in the absence of antibiotics.** For each experiment, replicate and strain, the table reports the rate at which tracked lineages are lost during the 30 min immediately before the switch antibiotic-containing media. Valid lineages are tracked lineages present at the start of the 30 min window. Lost lineages are valid lineages whose last valid frame falls inside the 30 min window. This is the same termination event that triggers the pseudo-area filling in Fig. 3 & 5. The loss rate (%) is the fraction of lineages present at the start of the window that are lost within these 30 antibiotic free minutes.

| **Exp.** | **Replicate** | **Strain** | **Valid lineages** | **Number of lost lineages** | **Loss rate (%)** |
|---|---|---|---|---|---|
| CTA 1 | 1 | EL4668 | 296 | 2 | 0.7 |
| | | EL4483 | 301 | 4 | 1.3 |
| | 2 | EL4668 | 313 | 1 | 0.3 |
| | | EL4483 | 310 | 4 | 1.3 |
| | 3 | EL4668 | 317 | 0 | 0.0 |
| | | EL4483 | 317 | 4 | 1.3 |
| CIP 0.25 → CTA 1 | 1 | EL4668 | 285 | 0 | 0.0 |
| | | EL4483 | 272 | 20 | 7.4 |
| | 2 | EL4668 | 314 | 1 | 0.3 |
| | | EL4483 | 299 | 11 | 3.7 |
| | 3 | EL4668 | 294 | 2 | 0.7 |
| | | EL4483 | 264 | 11 | 4.2 |
| MER 2 | 1 | EL4992 | 290 | 1 | 0.3 |
| | | EL4993 | 318 | 2 | 0.6 |
| | 2 | EL4992 | 287 | 0 | 0.0 |
| | | EL4993 | 280 | 0 | 0.0 |
| | 3 | EL4992 | 300 | 0 | 0.0 |
| | | EL4993 | 303 | 0 | 0.0 |
| MER 2 | 1 | EL4650 | 158 | 3 | 1.9 |
| | | EL4649 | 290 | 8 | 2.8 |
| | 2 | EL4650 | 208 | 0 | 0.0 |
| | | EL4649 | 293 | 8 | 2.7 |
| | 3 | EL4650 | 498 | 0 | 0.0 |
| | | EL4649 | 266 | 16 | 6.0 |
| MER 2 | 1 | EL4991 | 298 | 2 | 0.7 |

| | | | | | |
|---|---|---|---|---|---|
| | | EL4990 | 293 | 4 | 1.4 |
| | 2 | EL4991 | 299 | 0 | 0.0 |
| | | EL4990 | 299 | 5 | 1.7 |
| | 3 | EL4991 | 314 | 5 | 1.6 |
| | | EL4990 | 313 | 5 | 1.6 |
| MER 2 | 1 | EL5106 | 234 | 10 | 4.3 |
| | | EL4988 | 308 | 6 | 1.9 |
| | 2 | EL5106 | 210 | 17 | 8.1 |
| | | EL4988 | 261 | 2 | 0.8 |
| | 3 | EL5106 | 227 | 15 | 6.6 |
| | | EL4988 | 312 | 2 | 0.6 |
| CIP 0.25 | 1 | EL4900 | 286 | 2 | 0.7 |
| | | EL4884 | 297 | 3 | 1.0 |
| | 2 | EL4900 | 317 | 0 | 0.0 |
| | | EL4884 | 314 | 0 | 0.0 |
| | 3 | EL4900 | 311 | 2 | 0.6 |
| | | EL4884 | 307 | 3 | 1.0 |
| AMI 8 | 1 | EL4650 | 336 | 4 | 1.2 |
| | | EL4649 | 333 | 1 | 0.3 |
| | 2 | EL4650 | 307 | 0 | 0.0 |
| | | EL4649 | 317 | 0 | 0.0 |
| | 3 | EL4650 | 96 | 2 | 2.1 |
| | | EL4649 | 166 | 1 | 0.6 |
| CIP 0.25 | 1 | EL4668 | 302 | 3 | 1.0 |
| | | EL4824 | 291 | 5 | 1.7 |
| | 2 | EL4668 | 311 | 1 | 0.3 |
| | | EL4824 | 239 | 9 | 3.8 |
| | 3 | EL4668 | 303 | 1 | 0.3 |
| | | EL4824 | 282 | 0 | 0.0 |
| AMI 4 | 1 | EL4987 | 303 | 0 | 0.0 |
| | | EL4876 | 315 | 1 | 0.3 |
| | 2 | EL4987 | 305 | 0 | 0.0 |

| | | | | | |
|---|---|---|---|---|---|
| | | EL4876 | 293 | 3 | 1.0 |
| | 3 | EL4987 | 302 | 1 | 0.3 |
| | | EL4876 | 302 | 1 | 0.3 |
| CIP 0.25 → AMI 4 | 1 | EL4987 | 278 | 2 | 0.7 |
| | | EL4876 | 299 | 1 | 0.3 |
| | 2 | EL4987 | 314 | 1 | 0.3 |
| | | EL4876 | 313 | 2 | 0.6 |
| | 3 | EL4987 | 311 | 1 | 0.3 |
| | | EL4876 | 312 | 1 | 0.3 |
| CIP 1 | 1 | EL4821 | 260 | 0 | 0.0 |
| | | EL4670 | 220 | 5 | 2.3 |
| | 2 | EL4821 | 457 | 1 | 0.2 |
| | | EL4670 | 608 | 5 | 0.8 |
| | 3 | EL4821 | 321 | 5 | 1.6 |
| | | EL4670 | 579 | 8 | 1.4 |
| CIP 0.25 | 1 | EL4990 | 441 | 4 | 0.9 |
| | | EL4991 | 430 | 2 | 0.5 |
| | 2 | EL4990 | 358 | 11 | 3.1 |
| | | EL4991 | 402 | 0 | 0.0 |
| | 3 | EL4990 | 409 | 6 | 1.5 |
| | | EL4991 | 112 | 1 | 0.9 |
| AMI 8 | 1 | EL4871 | 302 | 0 | 0.0 |
| | | EL4880 | 311 | 0 | 0.0 |
| AMI 8 | 1 | EL4824 | 292 | 6 | 2.1 |
| AMI 8 | 1 | EL4987 | 310 | 2 | 0.6 |
| | | EL4876 | 309 | 0 | 0.0 |
| AMI 4 | 1 | EL4871 | 308 | 0 | 0.0 |
| | | EL4880 | 308 | 1 | 0.3 |
| AMI 4 | 1 | EL4824 | 289 | 5 | 1.7 |
| **Average loss rate** | | | | | **1.2** |

**Supplementary Table S3. Classification of resistant versus susceptible lineages from single-cell relative growth rate and relative area trajectories.** For each species and antibiotic condition, single lineage trajectories over time or generation normalized time, tau, were classified using PCA (two components) followed by logistic regression. One replicate was held out as a pure test set and the remaining replicates used for training. The replicates shown in the main figures, Figures 2 and 3, were always in the training set. The classifier was fit on the post-antibiotic window, with the pre-antibiotic window analysed identically as a control. Resistant lineages were defined as the positive class and all metrics are computed on the test replicate. $Window_{pre}$/$Window_{post}$ are the time/tau intervals used, $N_{training}$/$N_{test}$ are the lineages in the training and test sets, $AUC_{pre}$/$AUC_{post}$ are the ROC AUC for each window, while sensitivity/specificity are fractions of resistant/susceptible lineages correctly identified.

| **Species** | **Condition (μg/mL)** | **$Window_{pre}$** | **$Window_{post}$** | **$N_{training}$** | **$N_{test}$** | **$AUC_{pre}$** | **$AUC_{post}$** | **Sens.** | **Spec.** |
|---|---|---|---|---|---|---|---|---|---|
| *E. coli* | CIP 0.25 | [-30, 0] min | [0, 45] min | 1090 (R: 595, S: 495) | 581 (R: 307, S: 274) | 0.353 | 1.000 | 0.997 | 0.993 |
| *P. aeruginosa* | AMI 8 | [-30, 0] min | [0, 35] min | 1098 (R: 570, S: 528) | 208 (R: 89, S: 119) | 0.574 | 0.984 | 1.000 | 0.975 |
| *E. coli* | CIP 0.25 | [-30, 0] min | [0, 45] min | 1019 (R: 553, S: 466) | 546 (R: 294, S: 252) | 0.628 | 0.999 | 0.997 | 0.929 |
| *E. coli* | AMI 4 | [-30, 0] min | [0, 60] min | 1135 (R: 583, S: 552) | 554 (R: 301, S: 253) | 0.366 | 1.000 | 1.000 | 0.992 |
| *A. baumannii* | CIP 1 | [-30, 0] min | [0, 60] min | 963 (R: 478, S: 485) | 878 (R: 504, S: 374) | 0.548 | 0.990 | 0.982 | 0.968 |
| *Klebsiella* spp. | CIP 0.25 | [-30, 0] min | [0, 45] min | 1272 (R: 523, S: 749) | 672 (R: 384, S: 288) | 0.572 | 0.992 | 0.987 | 0.997 |
| *S. aureus* | CXI 4 | [-30, 0] min | [0, 60] min | 364 (R: 52, S: 312) | 374 (R: 134, S: 240) | 0.509 | 0.993 | 0.985 | 0.917 |
| *A. baumannii* | MER 2 | tau [-2, 0] | tau [0, 3] | 1057 (R: 521, S: 536) | 532 (R: 259, S: 273) | 0.647 | 0.994 | 0.981 | 0.978 |
| *E. coli* | CTA 1 | tau [-2, 0] | tau [0, 3] | 966 (R: 548, S: 418) | 480 (R: 282, S: 198) | 0.690 | 0.917 | 0.876 | 0.975 |
| *Klebsiella* spp. | MER 2 | tau [-2, 0] | tau [0, 4] | 965 (R: 534, S: 431) | 513 (R: 278, S: 235) | 0.576 | 0.983 | 0.968 | 0.928 |

**Supplementary Table S4. Per-species and per-isolate classification performance for the multi-species classifier.** Lineages from all experiments of a given condition were pooled and classified by PCA followed by logistic regression and evaluated by 5-fold cross-validation. All metrics are computed on the out-of-fold predictions. The condition gives the antibiotic and concentration used. For each species, N is the number of lineages. AUC, sensitivity and specificity are reported with resistant as the positive class. Within each species, individual isolates are listed by their MIC and n is the number of lineages for that isolate. The correct fraction is the fraction of n lineages classified correctly.

| **Condition (µg/mL)** | **Species** | **N** | **AUC** | **Sens.** | **Spec.** | **Isolate** | **n** | **Correct (frac)** |
|---|---|---|---|---|---|---|---|---|
| AMI 8 | *E. coli* | 1419 | 0.883 | 0.842 | 0.728 | MIC 1 | 290 | 288/290 |
| | | | | | | MIC 2 | 256 | 215/256 |
| | | | | | | MIC 8 | 289 | 105/289 |
| | | | | | | MIC 32 | 287 | 196/287 |
| | | | | | | MIC > 128 | 297 | 296/297 |
| | *P. aeruginosa* | 1279 | 0.995 | 0.982 | 0.994 | MIC 1 | 625 | 621/625 |
| | | | | | | MIC > 32 | 654 | 642/654 |
| MER 2 | *E. coli* | 870 | 0.957 | 0.883 | 0.906 | MIC 0.03 | 631 | 572/631 |
| | | | | | | MIC 16 | 239 | 211/239 |
| | *P. aeruginosa* | 709 | 0.767 | 0.472 | 0.997 | MIC 0.5 | 582 | 580/582 |
| | | | | | | MIC 16 | 127 | 60/127 |
| | *Klebsiella* spp. | 1547 | 0.924 | 0.965 | 0.694 | MIC 0.03 | 695 | 482/695 |
| | | | | | | MIC > 16 | 852 | 822/852 |
| | *A. baumannii* | 1608 | 0.997 | 0.991 | 0.943 | MIC 0.12 | 810 | 764/810 |
| | | | | | | MIC > 32 | 798 | 791/798 |

**Supplementary Table S5. Sequential classification of the same single-cell lineages across two consecutive antibiotic exposures.** Lineages from the sequence experiments were tracked continuously through a first CIP exposure and a second exposure to either AMI, classified from single-cell relative growth rate trajectories over time, or CTA, classified from relative cell area trajectories over generation normalized time, tau. The sequence replicates were held out as a test set and the replicates used for training, the replicates shown in Figures 2 and 3 were always in the training set. Resistant lineages were defined as the positive class. Because all strains in the test sets are CIP-resistant, the first stage only reports CIP-sensitivity, which is the fraction of test lineages correctly classified as resistant by the CIP classifier. The second stage performs the resistant versus susceptible classification reported by AUC, sensitivity and specificity. Condition (µg/mL) gives the two antibiotics applied in sequence with their concentrations. $N_{training}$/$N_{test}$ are the pooled lineage counts in the training and test sets. AUC, sensitivity and specificity are the ROC AUC, fraction of resistant lineages correctly identified and fraction of susceptible lineages correctly identified of the second stage classifier all computed on the test set.

| **Species** | **Condition (µg/mL)** | **$N_{training}$** | **$N_{test}$** | **Sens.$_{CIP}$** | **AUC** | **Sens.** | **Spec.** |
|---|---|---|---|---|---|---|---|
| *E. coli* | CIP 0.25 → AMI 4 | CIP: 3254, AMI: 1709 | 1667 | 0.992 | 0.998 | 0.987 | 0.989 |
| *E. coli* | CIP 0.25 → CTA 1 | CIP: 3254, CTA: 1540 | 1155 | 0.979 | 0.917 | 0.899 | 0.927 |

**Supplementary Table S6. Data from Figure 4.** For strains included in the figure, the table displays relative growth rates at AMI concentrations of 4 µg/mL or 8 µg/mL, measured at 30, 60 and 90 minutes post-AMI introduction. Each strain's relevant MIC (µg/mL) is indicated. N denotes the number of data points. Values shown are the mean relative growth rate with 25th and 75th percentiles.

| **Relevant MIC (µg/mL)** | **Time post-AMI** | **AMI concentration (µg/mL)** | **N** | **Mean of rel. growth rate (25th percentile, 75th percentile)** |
|---|---|---|---|---|
| 2 | 30 | 4 | 294 | 0.842 (0.805, 0.957) |
| | | 8 | 282 | 0.626 (0.488, 0.812) |
| | 60 | 4 | 289 | 0.225 (0.056, 0.357) |
| | | 8 | 273 | 0.035 (-0.005, 0.071) |
| | 90 | 4 | 281 | 0.001 (-0.052, 0.031) |
| | | 8 | 263 | -0.026 (-0.057, 0.004) |
| 8 | 30 | 4 | 300 | 0.975 (0.948, 1.023) |
| | | 8 | 309 | 0.874 (0.804, 0.999) |
| | 60 | 4 | 292 | 0.778 (0.669, 0.938) |
| | | 8 | 298 | 0.121 (-0.019, 0.243) |
| | 90 | 4 | 279 | 0.515 (0.219, 0.816) |
| | | 8 | 285 | -0.023 (-0.067, 0.005) |
| 32 | 30 | 4 | 303 | 0.957 (0.987, 1.033) |
| | | 8 | 303 | 0.863 (0.788, 1.008) |
| | 60 | 4 | 294 | 0.841 (0.790, 0.991) |
| | | 8 | 296 | 0.331 (0.002, 0.668) |
| | 90 | 4 | 287 | 0.661 (0.403, 0.956) |
| | | 8 | 281 | 0.200 (-0.040, 0.426) |
| > 128 | 30 | 4 | 308 | 1.054 (1.004, 1.109) |
| | | 8 | 302 | 1.008 (0.961, 1.039) |
| | 60 | 4 | 309 | 1.023 (0.977, 1.078) |
| | | 8 | 302 | 1.009 (0.962, 1.046) |
| | 90 | 4 | 308 | 1.039 (0.985, 1.095) |
| | | 8 | 302 | 1.012 (0.966, 1.039) |

### Supplementary Figures:

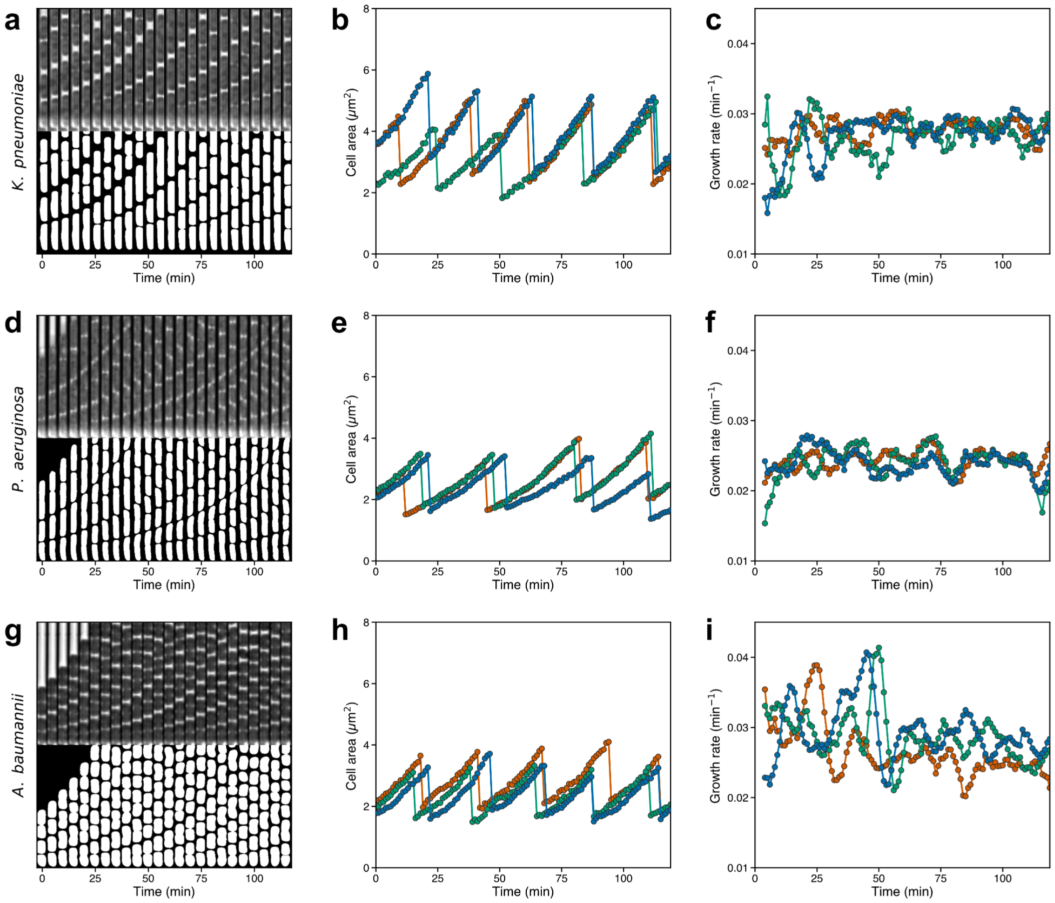


**Supplementary Figure S1:** As Figure 1 a-c, but for *K. pneumoniae*, (a-c) *P. aeruginosa*, (d-f) *A. baumannii (g-i)*.

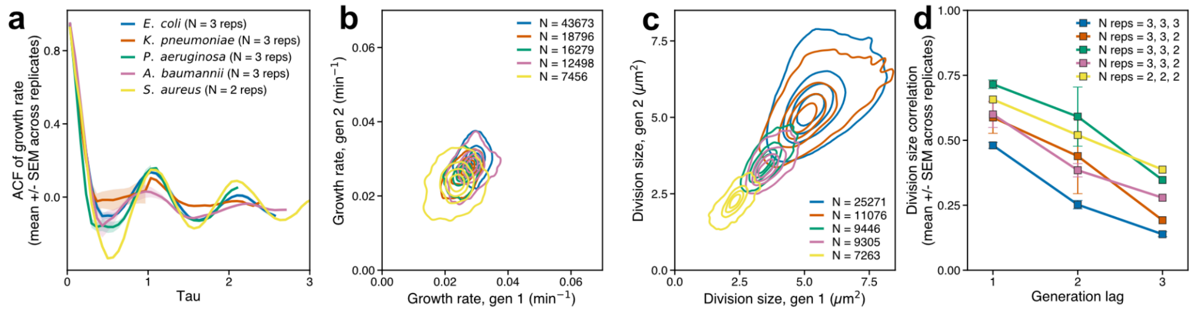


***Supplementary Figure S2:*** **Single-cell growth and division statistics.** Data shown for five bacterial species as indicated by marker color (**a**) Autocorrelation function (ACF) of single lineage growth rates as a function of dimensionless lag tau = t / <T_gen>, where <T_gen> average generation time during the period before antibiotic introduction. The ACF was computed per trap up to a maximum lag of N/2, where N is the number of frames in the time series, yielding 30-60 pairs at the maximum lag depending on series length. ACF curves were aggregated within each replicate by binning tau onto a common grid (1/15 generation bins), then averaged across the three replicates. The line shows the mean across replicates wherever at least one replicate has data; the shaded band shows the SEM across replicates and is shown only where all three replicates contribute. (**b**) Density of mother-daughter growth rate pairs, pooled across all replicates per species. (**c**) Density of mother-daughter division-size pairs (generation lag = 1 in panel (d)), pooled across all replicates per species. (**d**) Pearson correlation between division sizes at generation 1 and generation 1 + lag, computed per replicate; markers show the mean across replicates, error bars show the SEM. Numbers in the legends indicate the number of mother-daughter pairs (b, c) or replicates contributing at each lag (a, d). Pearson correlations are computed only where >= 3 mother-daughter pairs are available within a replicate.

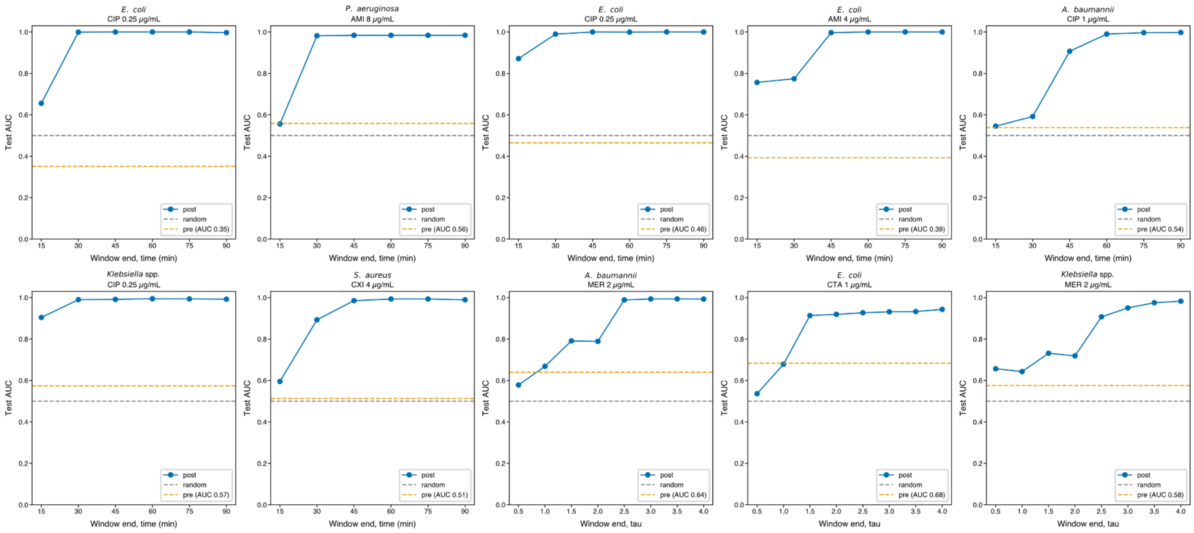


**Supplementary Figure S3:** AUC as function of time after start of antibiotic treatment (blue filled circles) for the isolate combinations used in Figure 2 and 3. Here AUCs are areas under the true vs. false resistance curves. Dashed black line is at 0.5 to highlight a random classification. Yellow dashed line is the result of performing susceptibility classification before antibiotic treatment.

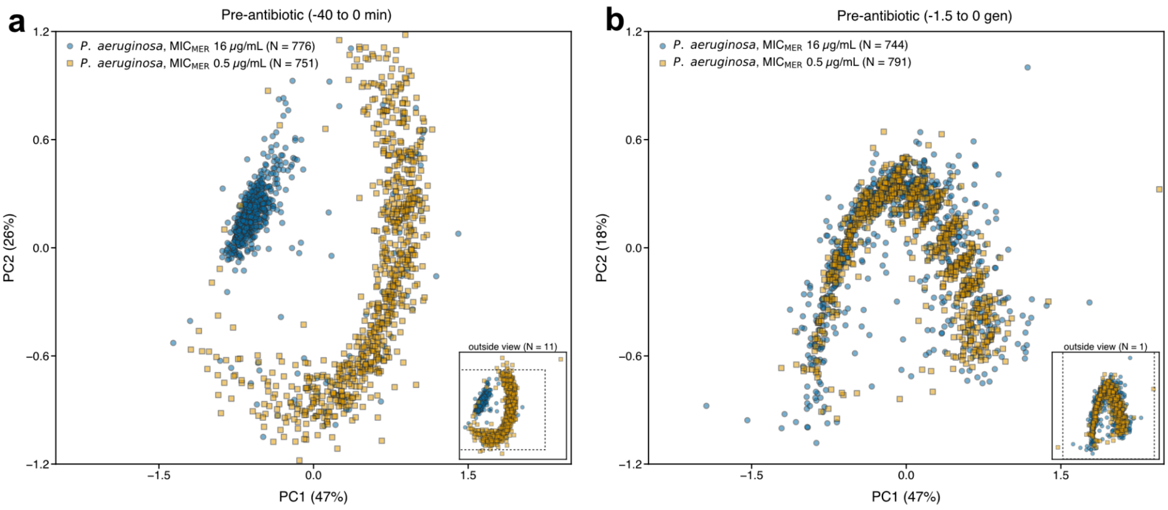


**Supplementary Figure S4:** (a) PCA of growth and division trajectories of two isolates of *P. aeruginosa* before normalizing time to the pre-treatment generation times for each cell lineage. (b) PCA after normalization using the pre-treatment generation time for each cell lineage.

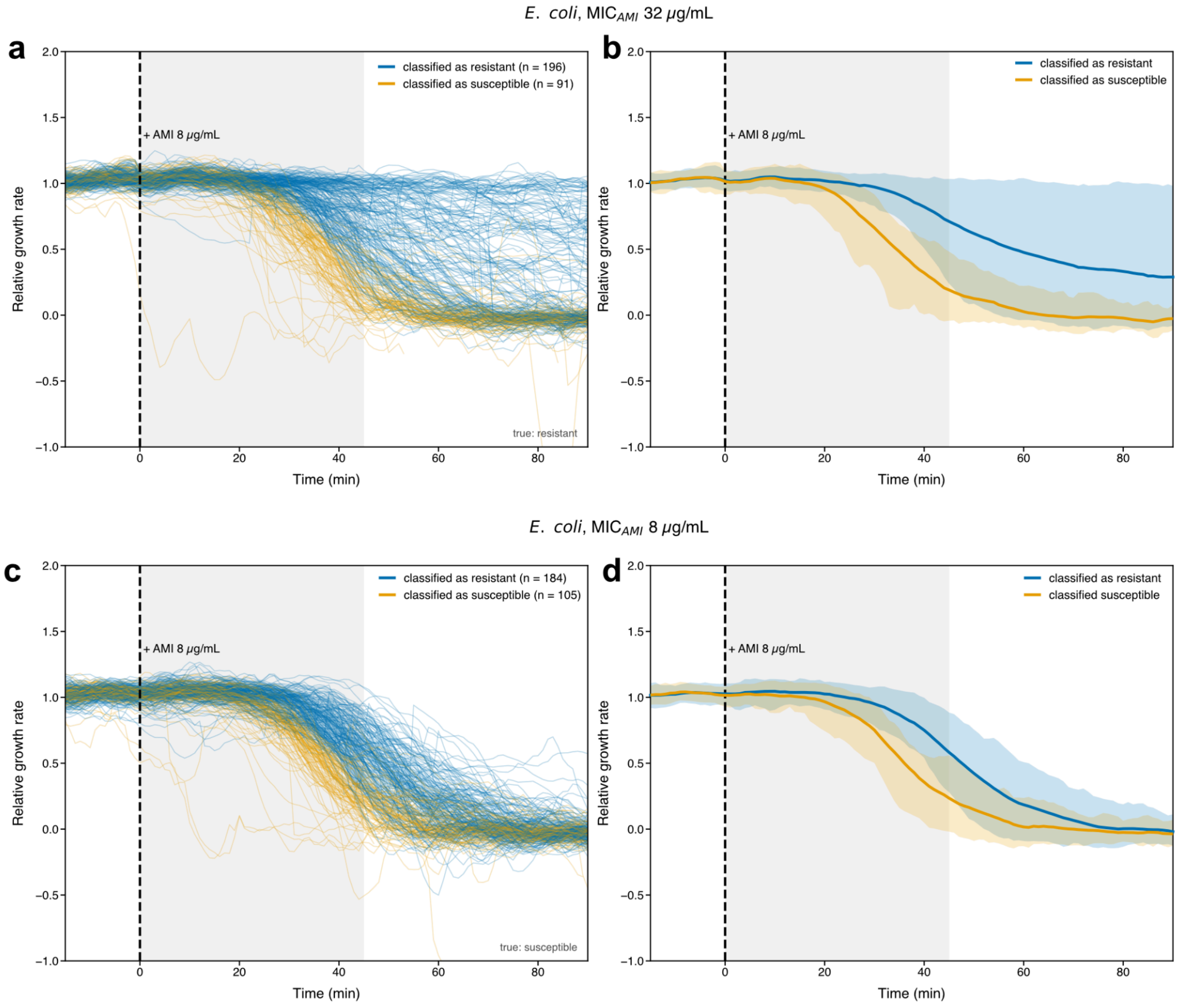


**Supplementary Figure S5:** (a & c) Relative growth rates of all cell lineages of the MIC$_{AMI}$ 8µg/mL (a) and MIC$_{AMI}$ 32µg/mL (c) isolates, both treated with 8µg/mL of AMI. Blue and orange curves are cell lineages classified as either resistant, or susceptible respectively in the analysis in Fig. 5 of the main text. (b & d) Average and 5% and 95% quantiles (as in Figure 2) of the cells in (a) and (c). Shaded regions indicate the time-window used for the classifier in Fig. 5.
